# Continuous lifelong learning for modeling of gene regulation from single cell multiome data by leveraging atlas-scale external data

**DOI:** 10.1101/2023.08.01.551575

**Authors:** Qiuyue Yuan, Zhana Duren

## Abstract

Accurate context-specific Gene Regulatory Networks (GRNs) inference from genomics data is a crucial task in computational biology. However, existing methods face limitations, such as reliance on gene expression data alone, lower resolution from bulk data, and data scarcity for specific cellular systems. Despite recent technological advancements, including single-cell sequencing and the integration of ATAC-seq and RNA-seq data, learning such complex mechanisms from limited independent data points still presents a daunting challenge, impeding GRN inference accuracy. To overcome this challenge, we present LINGER (LIfelong neural Network for GEne Regulation), a novel deep learning-based method to infer GRNs from single-cell multiome data with paired gene expression and chromatin accessibility data from the same cell. LINGER incorporates both 1) atlas-scale external bulk data across diverse cellular contexts and 2) the knowledge of transcription factor (TF) motif matching to *cis*-regulatory elements as a manifold regularization to address the challenge of limited data and extensive parameter space in GRN inference. Our results demonstrate that LINGER achieves 2-3 fold higher accuracy over existing methods. LINGER reveals a complex regulatory landscape of genome-wide association studies, enabling enhanced interpretation of disease-associated variants and genes. Additionally, following the GRN inference from a reference sc-multiome data, LINGER allows for the estimation of TF activity solely from bulk or single-cell gene expression data, leveraging the abundance of available gene expression data to identify driver regulators from case-control studies. Overall, LINGER provides a comprehensive tool for robust gene regulation inference from genomics data, empowering deeper insights into cellular mechanisms.

## Introduction

Gene regulatory networks (GRNs) [1] are collections of molecular regulators that interact with each other and determine gene activation and silencing in specific cellular contexts. A comprehensive understanding of gene regulation dynamics is fundamental to explain how cells perform diverse functions despite sharing the same genome, how cells alter gene expression in response to different environments, and how noncoding genetic variants cause disease. GRNs are composed of transcription factors (TFs) that bind DNA regulatory elements (REs) to activate or repress the expression of target genes (TGs).

Inference of GRNs from experimental data is a central problem [1,2], and there have been many attempts to approach this problem [1,3–9]. Co-expression-based methods such as WGCNA [10], ARACNe [7], and GEINE3 [11] infer the {TF-TG} *trans*-regulation from gene expression data by capturing the TF-TG covariation across multiple conditions/samples through linear or nonlinear methods. Such networks have undirected edges, preventing distinction between TFA regulating TFB or vice versa from a TF_A_–TF_B_ edge. Moreover, co-expression networks edges are interpreted as correlations between gene pairs rather than causal regulation. Genome-wide measurements of chromatin accessibility, such as DNase-seq [12] and ATAC-seq [13], provide rich information on mechanisms of gene regulation because open chromatin regions typically have less compacted DNA and are more accessible to transcription factors. Chromatin accessibility data provides the location of REs enabling motif matching to connect TF-RE and connecting REs to their nearby TGs so that we can infer the TF-TG relationship [14]. However, TF footprint approaches cannot distinguish within-family TFs sharing motifs. To overcome this limitation, we developed a statistical model, PECA [15], to fit TG expression by TF expression and RE accessibility from paired gene expression and chromatin accessibility data across a diverse panel of cell types. However, the problem still has not been fully solved because heterogeneity of cell types in bulk data limits the accuracy of inference. Bulk data are aggregated over millions of cells, and most biological samples are a mixture of cell types. Thus, GRN inference from bulk data may be not relevant to any included cell types.

The advent of single-cell (sc) sequencing technology has enabled highly accurate investigation of regulatory analysis at the level of individual cell types. Single-cell RNA sequencing (scRNA-seq) data, enables cell type-specific *trans*-regulation inference through co-expression analysis such as PIDC and SCENIC [16–24]. Single-cell chromatin accessibility sequencing (scATAC-seq) can be used to infer *trans*-regulation, such as DeepTFni [25]. Many methods integrate unpaired scRNA-seq and scATAC-seq data, which are measured not on the same cell but on two batches of the same mixed population, to infer *trans*-regulation. Those methods, including IReNA[26], SOMatic [27], UnpairReg [28], CoupledNMF [29,30], DC3 [30], and Wang et al. [31], link TFs to REs by motif matching and link REs to TGs using the covariation of RE-TG across cell types or physical base pair distance. Despite extensive efforts, GRN inference accuracy has remained disappointingly low, marginally exceeding random predictions [32].

Recent advances in sc sequencing as well as ATAC-seq and RNA-seq integration [33] provide opportunities to address these challenges effectively [34], exemplified by the new computational method SCENIC+ [35] in GRN inference. However, three major challenges persist in GRN inference. First, learning such a complex mechanism from limited data points remains a significant obstacle. Although single-cell data offers a large number of cells, most of them are not independent, restricting the number of distinct conditions available for analysis. Second, incorporating prior knowledge like TF-RE motif matching data into nonlinear models like neural networks is challenging. Third, inferred GRN accuracy assessed by experimental data such as ChIP-seq and perturbations is only marginally better than random prediction [32].

To overcome these challenges, we propose a novel method called LIfelong neural Network for GEne Regulation, referred to as LINGER. This research paper contributes to the field of GRN inference in multiple ways. First, LINGER employs lifelong learning to incorporate large-scale external bulk data and neural networks to model TG expression by TFs and REs, mitigating the challenge of limited data but extensive parameters to be estimated. Second, LINGER integrates TF-RE motif matching knowledge through manifold regularization term in the cost function, enabling prior knowledge incorporation into the modeling process. Third, it substantially improves GRN inference accuracy, as indicated by the significant increase in the ratio of Area Under the Precision-Recall Curve (AUPR) compared to a random predictor from the traditional value of 1.2 to 3. It is a substantial improvement because the previous methods achieve about only 20% more accurate prediction compared to a random predictor but LINGER achieves about 2-fold more accuracy than a random predictor. Fourth, based on the inferred cell type specific GRNs, LINGER enables the estimation of TF activity solely from gene expression data such as sc/bulk RNA-seq or microarray data, identifying driver regulators between disease/healthy groups. Fifth, as a testament to the openness of scientific research, the LINGER software is publicly available to the scientific community. Researchers can utilize this tool to enhance their own GRN inference studies and further contribute to the advancement of this field.

In conclusion, this research paper presents LINGER as an innovative approach to improve GRN inference from sc-multiome data. The incorporation of external data by lifelong learning, and prior knowledge integration by manifold regularization enables a significant enhancement in GRN accuracy. Furthermore, the ability to estimate TF activity solely from gene expression data offers valuable insights into disease mechanisms and potential therapeutic targets.

## Results

### LINGER: leveraging external bulk data to infer gene regulatory networks from single cell multiome data

LINGER is a computational framework designed to infer GRNs from single-cell multiome data (Fig.1 and Methods). Using count matrices of gene expression and chromatin accessibility, along with cell type annotation as input, it provides a population level general GRN, cell type specific GRNs, and cell-level GRNs. Each GRN contains three types of interactions among three types of nodes {TF, RE, TG}, namely, *trans*-regulation {TF-TG}, *cis*-regulation {RE-TG}, and TF-binding {TF-RE}. The {TF-TG}, {RE-TG}, and {TF-RE} relationships, which represent TF regulation of TG expression, *cis*-RE regulation of TG expression, and TF binding to *cis*-REs, respectively. LINGER is distinguished by its ability to integrate the comprehensive gene regulatory profile from external paired gene expression and chromatin accessibility bulk data across diverse panels of cellular contexts. This is achieved through a lifelong machine learning, also called continuous learning, based on neural network (NN) models, which has been proven to leverage the knowledge learned in previous tasks to help learn the new task better [36].

**Fig. 1.**
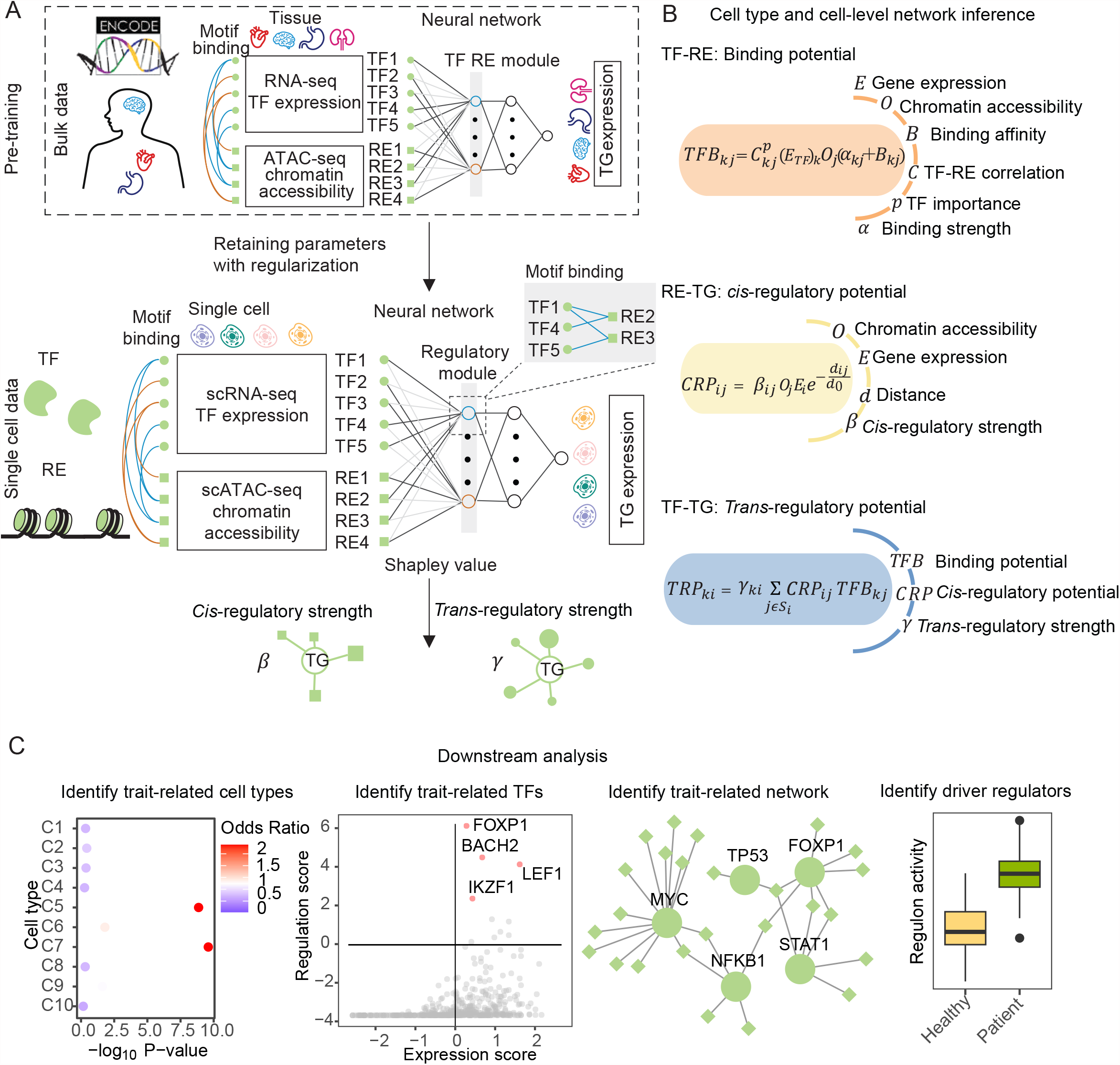
Schematic overview of LINGER. **A**. Schematic illustration of LINGER. LINGER is a model predicting gene expression by TF expression and chromatin accessibility using a deep neural network model. LINGER pretrains on the atlas-scale external bulk data and retains parameters via lifelong learning. The population-level GRN is generated from the neural network using the Shapley value. **B**. Strategy for constructing cell type specific and cell-level GRNs. Cell type specific and cell-level GRNs are inferred by the identical strategy, which combines consistent information across all cells, including regulatory strength, motif binding affinity, and RE-TG distance, with context-specific information on gene expression and chromatin accessibility. **C**. Downstream analyses enabled by LINGER-inferred GRNs, including identifying complex regulatory landscape of GWAS traits and driver regulator identification.

LINGER method is proposed to mitigate the challenge of limited independent data for inferring GRNs with a large number of parameters. LINGER leverages external bulk data to enhance the inference process from single-cell multiome data, incorporating three key steps: training on external bulk data, refining on single-cell data, and extracting regulatory information using interpretable AI techniques. In our approach, we utilize an NN model to fit the expression of TGs, taking as input of TF expression and the accessibility of REs within a 1 Mb distance from the transcription start site (TSS) of the TGs. The second layer of the NN model consists of weighted sums of TFs and REs, forming regulatory modules. The formation of these regulatory modules is guided by TF-RE motif matching, which leverages existing knowledge and simplifies the model. Specifically, we incorporate manifold regularization into the loss function based on TF-RE motif matching graphs. This leads to the enrichment of TF motifs binding to REs that belong to the same regulatory module. First, we commence the model training using external bulk data obtained from the ENCODE project [37], which contains hundreds of samples covering diverse cellular contexts.

For refinement on single-cell data, we initialize the model with the parameters obtained from the bulk data and introduce a cosine distance based elastic weight consolidation (EWC) loss to prevent significant deviation of parameter estimation from the original bulk data solution. The magnitude of parameter deviation from the bulk data solution is determined by the second-order derivatives (Fisher information) of the estimated parameters, which reflects the sensitivity of the loss function to parameter changes in bulk data. In the Bayesian context, we can view the prior knowledge gained from the bulk data as the prior distribution, which forms our initial beliefs about the model parameters. As the model undergoes training and learns from new single cell data, the posterior distribution is updated, combining the prior knowledge with the likelihood of the new data. The EWC loss can be interpreted as a regularization term that encourages the posterior distribution to remain close to the prior distribution. This way, the model retains its prior knowledge while adapting to the new data, preventing excessive changes and ensuring a more stable and continuous learning process [38]. After training the NN model on both bulk and single-cell data, we infer the regulatory strength of TF-TG and RE-TG interactions using the Shapley value of the NN models, which estimates the contribution of each feature. The TF-RE binding strength is generated by the correlation of TF and RE based on the parameter learned in the second layer of the NN model. This comprehensive approach enables the inference of GRNs by incorporating external bulk data, while maintaining interpretability using regulatory modules and motif matching (see Figure 1A). LINGER then constructs the cell type specific and cell-level GRNs based on the general GRN and the cell type specific profiles (Fig. 1B). We use the identical strategy to infer cell type specific and cell-level GRN, which combines consistent information across all cells, including regulatory strength, motif binding affinity, and RE-TG distance, with context-specific information of gene expression and chromatin accessibility (See “GRN inference by Lifelong learning” in Methods).

We will use independent datasets to validate the inference of GRN below, and then we perform several downstream analyses: first, identification of the disease/trait related cell type, TFs, and GRN combining Genome-wide association studies (GWAS) data; second, constructing regulon activity on external expression data based on the corresponding GRN and identifying driver regulators as differentially active TFs between diseased and control individuals (Fig. 1C).

### LINGER improves the cellular population GRN inference by lifelong machine learning

To access the performance of LINGER, we utilized a public multiome dataset of peripheral blood mononuclear cells (PBMCs) from 10X Genomics (see the “Methods” for detail). It has been believed that a regression-based GRN inference method could accurately predict gene expression, which is indicative of its capacity to capture the underlying regulatory mechanisms and yield improved GRN inference. To investigate whether a linear model is adequate for modeling gene expression or a non-linear model is necessary, we conducted a comparative analysis between two models. The first model employs an Elastic Net framework to predict the expression of TG by TFs and REs. The second model, referred to as scNN, was a three-layer deep neural network model. The scNN model, sharing LINGER’s architecture but without motif matching information and pretraining using external bulk data, was designed to incorporate non-linear interactions and capture complex regulatory patterns inherent in the gene expression data.

We assessed the gene expression prediction ability of two models using 5-fold cross-validation and measured the Pearson correlation coefficient (PCC) on the test sets. We found that the scNN could model the gene expression better than Elastic net with a −*log*_10_ p-value of 569.11 from a paired *t*-test, especially for unpredictable genes by Elastic Net (PCC<0) (−*log*_10_ p-value: 1095.50, Fig. 2A). This demonstrates that the three-layer deep neural network model scNN outperforms the Elastic Net model in predicting gene expression, highlighting the superior predictive capability of the non-linear model in capturing the complex regulatory patterns within the GRN.

**Fig. 2.**
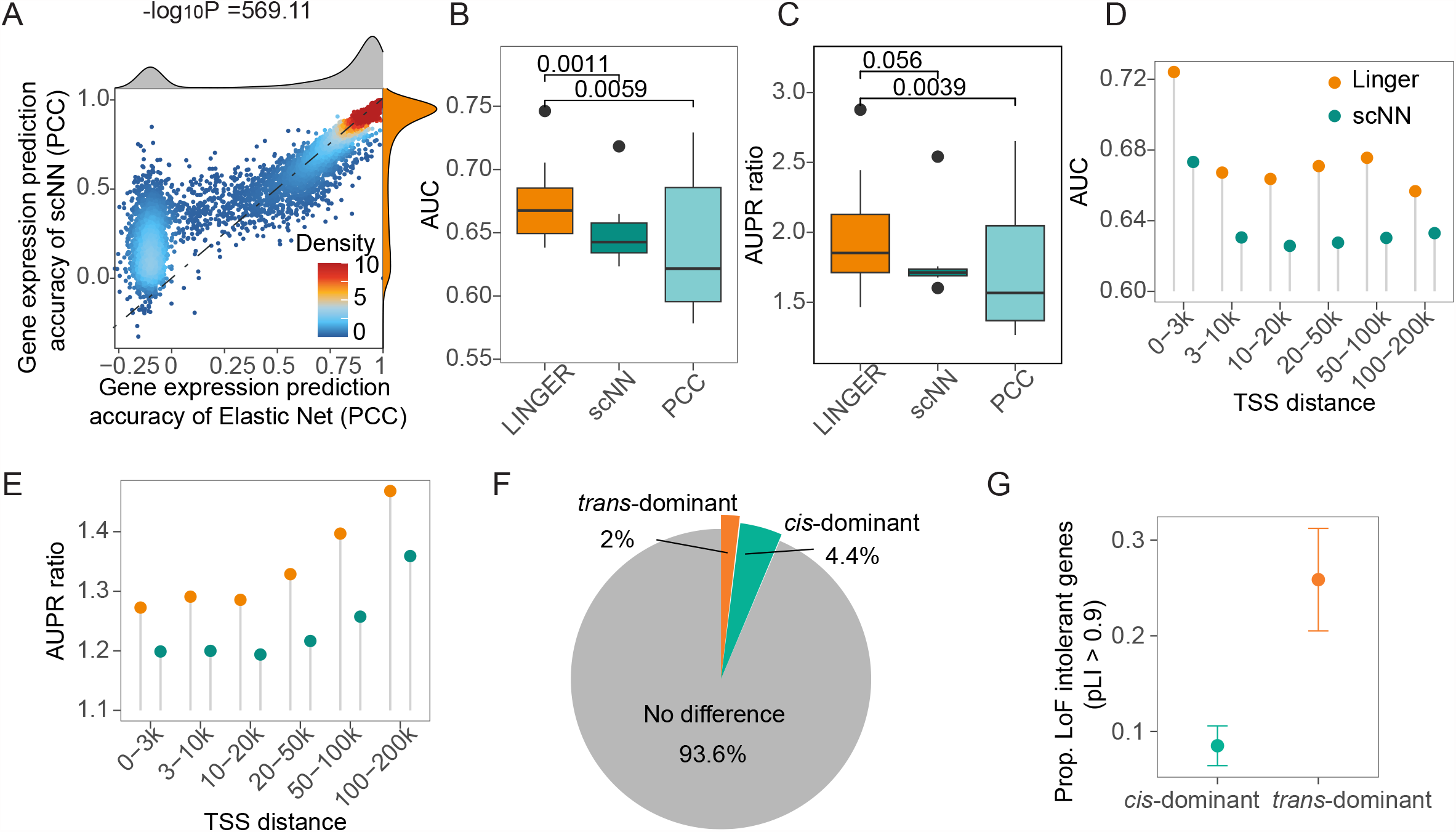
LINGER improves the cellular population GRN inference. **A**. Correlation between predicted and real gene expression, showing higher accuracy for LINGER versus ElasticNet. The x-axis represents the PCC of genes between ElasticNet predicted and real gene expression across cells, while the y-axis gives the PCC for LINGER. The points represent genes, and the color of the points represents the density. The color of distribution and B to E indicates the different methods. Orange represents LINGER, gray represents ElasticNet, dark green represents scNN, and light blue represents PCC. **B**. Boxplot of the performance metrics AUC for the predicted *trans*-regulatory strength across all ground truth data. The ground truth for B and C is putative targets of TFs from ten ChIP-seq data in blood. PCC denotes Pearson’s correlation coefficient between the chromatin accessibility of RE and the expression of TG. **C**. Boxplot of the performance metrics AUPR ratio for the predicted *trans*-regulatory strength. **D**. AUC for *cis*-regulatory strength inferred by LINGER. The ground truth for D and E is the variant-gene links from eQTLGen. We divide RE-TG pairs into different groups based on the distance of RE and the TSS of TG. **E**. AUPR ratio for *cis*-regulatory strength. **F**. Classification of the *trans*-dominant or *cis*-dominant gene. TFs contribute more to predicting the expression of *trans*-dominant genes, while REs contribute more to *cis*-dominant genes. **G**. Probability of *trans*-dominant and *cis*-dominant to be LoF intolerant genes. *Trans*-dominant genes are more likely to be loss-of-function intolerant.

To show the utility and effectiveness of integrating external bulk data, we conducted a comparative analysis of the GRN inference capabilities of LINGER in relation to the baseline model scNN, as well as other existing methods. To evaluate the performance of *trans*-regulatory strength, we collected putative targets of TFs from ChIP-seq data, and in total, we got 10 data in blood as ground truth [39](Table. S1). For each ground truth, we calculated the area under the receiver operating characteristic (ROC) curve (AUC) and the area under the precision-recall (PR) curve (AUPR) ratio (see the “Methods” for detail) by sliding the *trans*-regulatory predictions. Results show scNN performs better than PCC of TF-TG pairs. Compared to PCC and scNN, LINGER performs better with significantly higher AUC (P = 1.1e-3 for scNN and P = 5.9e-3 for PCC, paired *t*-test; Fig. 2B) and AUPR ratio (P = 3.9e-3 for PCC and P = 0.056 for scNN, which is a borderline significance; Fig. 2C) across all ground truth data.

To validate the *cis*-regulatory inference of LINGER, we calculated the consistency of the *cis*-regulatory coefficients with expression quantitative trait loci (eQTL) studies that link genotype variants to their target genes. We downloaded variant-gene links defined by eQTL in whole blood from GTEx [40] and eQTLGen [41] as ground truth. As the distance between RE and TG is important for the prediction, we divided RE-TG pairs into different distance groups (0-3 kb, 3-10 kb, 10-20 kb, 20-50 kb, 50-100 kb, and 100-200 kb). In each distance group, we calculated AUC and AUPR, with LINGER achieving higher AUC and AUPR than scNN in all different distance groups in eQTLGen (Fig. 2D and E) as well as GTEx (Fig. S1A and B). The above results show that LINGER improves the *cis*- and *trans*-regulatory strength inference by leveraging external data.

We next sought to investigate the dominant regulation for genes, that is, a gene is mainly regulated by *cis* or *trans*-regulation. To shed light on this question, we compared the average of *cis*- and *trans*-regulatory strength, the Shapely values of input features, via an unpaired t-test and perform Bonferroni p-value correction. Our findings reveal that the majority of genes exhibit no significant difference in *cis* and *trans*-regulation dominance. Specifically, 4.37% of genes are *cis*-regulation dominant, while 2.00% are *trans*-regulation dominant (Fig. 2F).. To discern evolutionary distinctions between *trans*-dominant and *cis*-dominant genes, we compared their strength of selection using pLI, which is an estimate of the “probability of being loss of function intolerant” [42]. We observed that the percentage of selectively constrained genes with high pLI (>0.9) in the *trans*-dominant group was approximately three times higher than that in the *cis*-dominant group (Fig. 2G). A previous study found that disease-associated genes from GWAS were enriched in selectively constrained genes, while eQTL genes were depleted in selectively constrained genes [43]. These observations highlight the importance of the *trans*-regulatory network in understanding complex diseases. Functional enrichment analysis [44] shows that the *cis*-regulatory dominant genes were significantly enriched in 38 GTEx aging signatures (Table. S2).

### LINGER improves the cell type specific GRN inference

We evaluated the cell type specific GRN inference of LINGER in PBMCs sc-multiome data as well as an in-silico mixture of H1, BJ, GM12878, and K562 cell lines from droplet-based single-nucleus chromatin accessibility and mRNA expression sequencing (SNARE-seq) [45] data. To evaluate the performance of our method, we established a reliable ground truth for each type of GRN. To assess TF-RE binding prediction, we utilized ChIP-seq data as ground truth, including ten TFs from four cell types within the blood and 33 TFs from the H1 cell line [39] (Table. S1). The putative target of TF from the ChIP-seq data serves as ground truth for inference of *trans*-regulatory potential. For *cis*-regulatory potential, we incorporated promoter capture HiC data of three primary blood cell types [46] and single-cell eQTL [47] including six immune cell types as ground truth for PBMCs.

The cell type-specific TF-RE binding potential in LINGER is determined by five factors: 1) cell type specific TF expression, 2) cell type specific RE chromatin accessibility, 3) motif binding affinity, 4) TF-RE binding strength, and 5) TF-RE correlation weighted by TF importance score in the cell type (see “Construct cell type-specific gene regulatory network” in Methods). We compared our method with TF-RE correlation (PCC) and motif binding affinity. To assess the TF-RE binding potential, we generated the ROC curve and PR curve for each ground truth data and then calculated the AUC and AUPR ratio, respectively. For example, in naive CD4 T cells, LINGER achieves an AUC of 0.92 and an AUPR ratio of 5.17 for ETS1, which is obviously better than PCC (AUC: 0.78, AUPR ratio: 2.71) and motif binding affinity (AUC: 0.70, AUPR ratio: 1.92) (Fig. 3A and E). For binding sites of MYC in the H1 cell line, LINGER (AUC: 0.84, AUPR ratio: 3.47) outperforms PCC (AUC: 0.42, AUPR ratio: 0.83) and motif binding affinity (AUC: 0.65, AUPR ratio: 1.51) based predictions (Fig. S2 A and B). For all 10 TFs in PBMC and 29 out of 33 TFs in H1, LINGER consistently exhibits highest AUC and AUPR ratios compared to the other two methods, and the overall distribution of AUC and AUPR ratios are significantly higher than others in both PBMC (P<=2.48e-3, Fig. 3B and C) and H1 data (P <= 6.68e-06, Fig. S2C and D) by one sided paired t-test. Furthermore, we compared LINGER with a state-of-the-art method, SCENIC+ [35], which predicts TF-RE pairs from multiome single-cell data. Since SCENIC+ does not provide a continuous score for all REs, it is not feasible to utilize metrics AUC and AUPR ratios for evaluating performance. We use *F*_1_ score as a measure of the accuracy by selecting the top TF-RE to generate the same number of TF-RE pairs as SCENIC+. Fig. 3D shows that LINGER performs better for all 10 TFs’ binding site prediction in PBMC data (paired t-test, p-value 0.02).

**Fig. 3.**
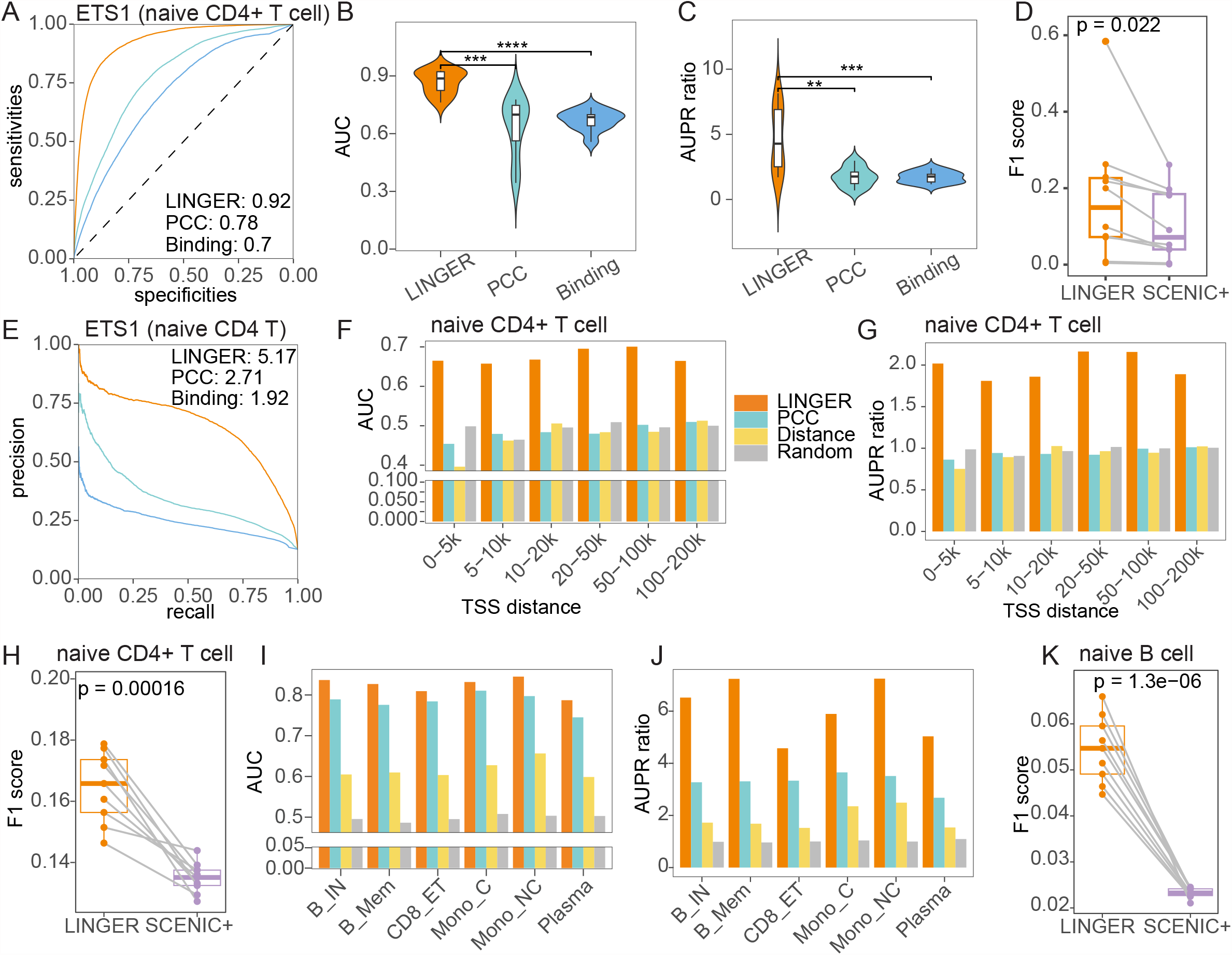
Systematic benchmarking of cell type specific TF-RE binding potential and *cis*-regulatory potential performance. **A, E**. ROC curve and PR curve of binding potential for ETS1 in naive CD4 T cells. The ground truth for A to E is the ChIP-seq data of ETS1 in naïve CD4+ T cells. The color in A to E represents the different competitors to predict TF-RE regulation. Orange represents LINGER, green represents PCC between the expression of TF and the chromatin accessibility of RE, and blue represents motif binding affinity of TF to RE. **B, C**. Violin plot of AUC and AUPR ratio values of binding potential across diverse TFs and cell types. The ground truth is ChIP-seq data for 10 TFs from different cell types in blood. In this study, we use the following convention for symbols indicating statistical significance: ns: p > 0.05; *: p <= 0.05; **: p <= 0.01; ***: p <= 0.001; ****, p <= 0.0001. We hide the ns symbol when displaying significance levels. **D**. The performance metrics F1 score of binding potential. Each point represents for each ground truth data. The p-value for D, H, and K are based on the one sided paired t-test. **F, G**. AUC and AUPR ratio of *cis*-regulatory potential in naïve CD4+ cell. The ground truth for F to H is promoter capture HiC data. RE-TG pairs are divided into six distance groups ranging from 0-5k to 100-200 kb. PCC is calculated between the expression of TG and the chromatin accessibility of RE. Distance denotes the decay function of the distance to the TSS. Random denotes the uniform distribution. **H**. F1 score of *cis*-regulatory in naïve CD4+ cell for LINGER and SCEINC+. **I, J**. AUC and AUPR ratio of *cis*-regulatory potential. The ground truth is eQTL data from 6 immune cell types. **K**. F1 score of *cis*-regulatory potential in naïve B cell. The ground truth is eQTL data from naïve B cells. This figure corresponds to the PMBC data.

To generate cell type specific *cis*-regulatory potential for RE-TG pairs, we combined several factors such as cell type specific TG expression, RE chromatin accessibility, *cis*-regulatory strength, as well as exponential decay of the genomic distance between RE and TG. We compared our method with four baseline methods including distance-based methods, PCC between enhancer and promoter accessibility, and random predictions. We also compared our method with SCENIC+, which outputs predicted RE-TG pairs without importance scores so that we use F1 score to compare it with LINGER. To evaluate the accuracy using HiC data, we divided RE-TG pairs into six distance groups ranging from 0-5k to 100-200 kb. In naïve CD4 T cells, LINGER achieves AUC ranging from 0.66 to 0.70 (Fig. 3F), and AUPR ratios ranging from 1.81 to 2.16(Fig. 3G) across all distance groups, while the performances of other methods are close to random. In other cell types, including naïve CD8 T cells and naïve B cells, LINGER exhibits consistent superiority over the baseline methods in terms of AUC and AUPR (Fig. S2E-H). As for eQTL data based validation, we regard all eQTL pairs as positive labels since there are not enough pairs to divide into distance groups. In all cell types, the AUC and AUPR ratios of LINGER are higher than the baseline methods (Fig. 3I and J). When comparing LINGER with SCENIC+, we selected the same number of top-ranking RE-TG pairs based on the *cis*-regulatory potential and calculated the F1 score using nine cutoffs corresponding to the 10th, 20th, …, and 90th percentile quantiles. As a result, LINGER attains significantly higher *F*_1_ scores than SCENIC+ in naïve CD4 T cells, naïve CD8 T cells, and naïve B cells (paired *t*-test p-values 1.6e-4, 2.7e-3, and 3.8e-3, respectively, Fig. 3H, Fig. S2I and J). Taking eQTL as ground truth, *F*_1_ score of LINGER is significantly higher than SCENIC+ under nine equal quantile cutoffs in naïve B cells (P=1.3e-6, paired *t*-test; Fig. 3K) and other cell types (Fig. S2 K-O).

Cell type specific *trans*-regulatory potential for TF-TG pairs considers all REs within 1Mb around the TSS of the TG. To represent the contribution of each RE to the TF-TG trans-regulatory potential, we multiply the RE-TG *cis*-regulatory potential with the TF-RE binding potential. The TF-TG score is then defined as the sum of the contributions from all the REs considered. To evaluate the accuracy of *trans*-regulatory potential, we chose GENIE3 [11] and PIDC [16] for comparison based on benchmarking literature of GRN inference from single-cell data [32] as we choose in previous work [48] (see ‘Benchmark the trans-regulatory potential’ in Methods). In addition, we compared LINGER with PCC and SCENIC+. For target identification of STAT1 in classical monocytes, LINGER obviously improves the performance of prediction, as evidenced by AUC of 0.76 versus 0.57-0.59 and AUPR ratio of 2.60 versus 1.26-1.36 (Fig. 4A and B). A similar improvement is observed for CTCF in H1, with an increase in AUC from 0.44-0.50 to 0.66 and increase in AUPR ratio from 0.84-1.00 to 1.61 (Fig. S2P and Q). Overall, LINGER consistently performs better than other methods both in all TFs in PBMCs and all TFs in the H1 cell line with significantly higher AUC and AUPR ratio (P<= 2.15e-3 Fig. 4C and D; Fig. S2R).

**Fig. 4.**
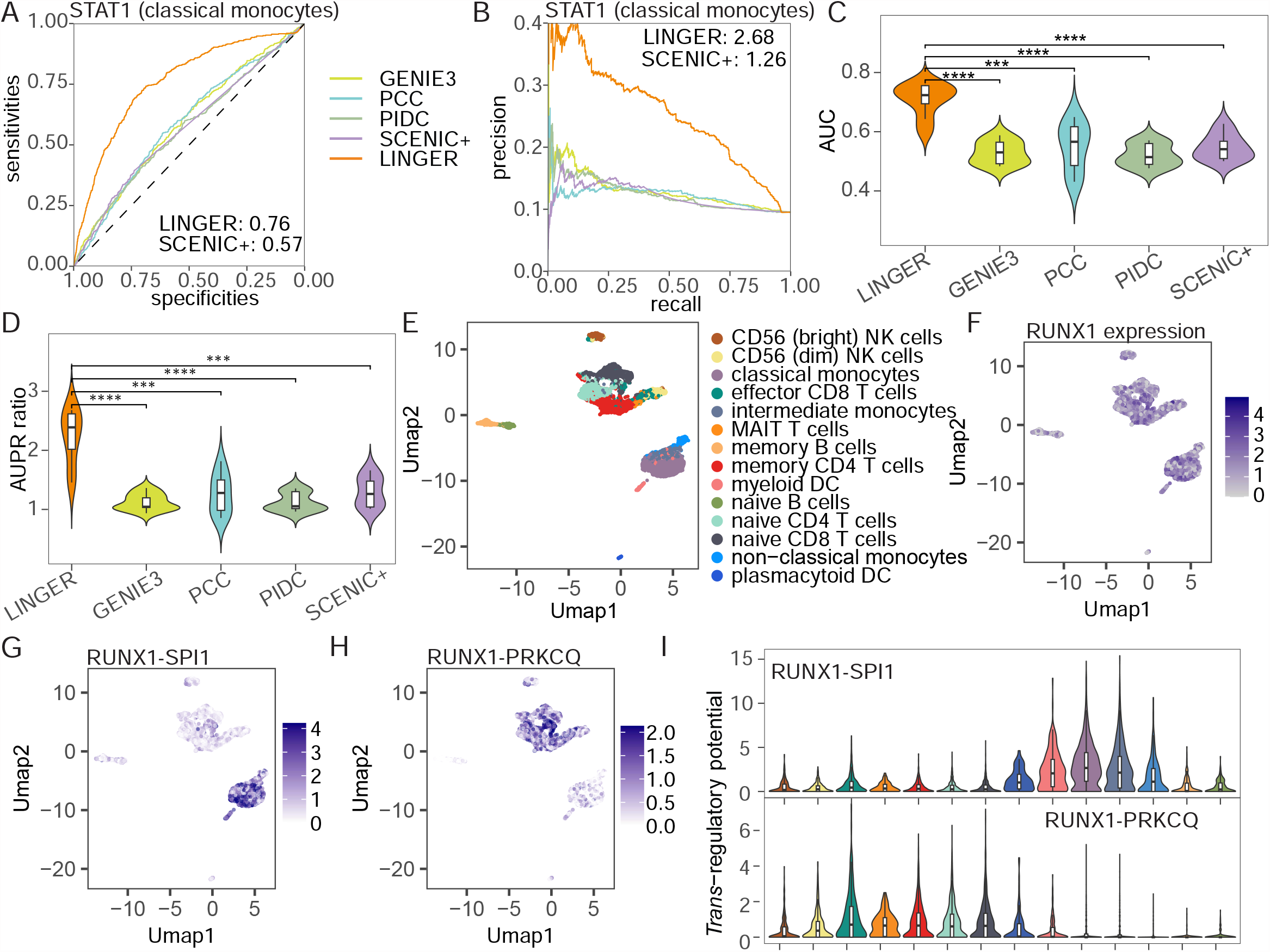
Systematic benchmarking of cell type specific *trans*-regulatory potential performance. **A, B**. ROC curve and PR curve of *trans*-regulatory potential inference of STAT1 in classical monocytes. The ground truth of A to D is putative targets of TFs from ChIP-seq data for the corresponding cell types in PBMCs. **C, D**. Violin plot of AUC and AUPR ratio values of *trans*-regulatory potential performance across diverse TFs and cell types. **E**. Umap of PBMCs including 14 cell types. **F**. Umap of RUNX1 expression across PBMCs. **G**. Umap of cell level *trans*-regulatory potential for RUNX1(TF)-SPI1(TG) across PBMCs. **H**. Umap of cell level *trans*-regulatory potential for RUNX1(TF)-PRKCQ(TG) across PBMCs. **I**. Violin plot of cell level *trans*-regulatory potential from different cell types. This figure corresponds to the PBMCs.

The rationale for constructing single-cell level GRN is the same with cell type specific GRN, replacing the cell type specific term with the single-cell term, such as gene expression and chromatin accessibility (see “Construct cell-level gene regulatory network” in Methods). We show the result of *trans*-regulation, taking RUNX1 as an example. RUNX1 is critical for the establishment of definitive hematopoiesis [49] and expresses at high levels in almost all PBMC cell types (Fig. 4E and F). RUNX1 regulates SPI1 in monocytes (classical, non-classical, and intermediate) and myeloid DC (Fig. 4G and I), while regulates PRKCQ in CD56 (dim) NK cells, effector CD8 T cells, MAIT T cells, memory CD4 T cells, naive CD4 T cells, and naive CD8 T cells (Fig. 4H and I). This example illustrates the capability of LINGER to visualize gene regulation at the single-cell level, offering valuable insights into cell type specific gene regulation. By visualizing gene regulation in individual cells, LINGER offers a powerful tool for unraveling the complexities of cell type-specific gene regulation and gaining valuable insights into cellular processes and functions.

### LINGER reveals the complex regulatory landscape of GWAS traits

Genome-wide association studies (GWASs) have successfully identified thousands of disease-associated variants, but the cells and functional programs in which genes are regulated by these variants are active remain largely unknown [50]. We propose a method to integrate GWAS summary statistic data and cell type specific GRN to identify the relevant cell types, key TFs, and sub-GRN implicated in those variants (see Methods). We define a trait regulation score for each TF in each cell type, measuring the enrichment of GWAS genes in downstream of the TF based on the cell type specific GRN. The trait regulation score should be independent of TF expression in non-relevant cell types. However, in trait-relevant cell types, a TF with a high trait regulation score should be expressed to perform its function effectively. We identify the trait-relevant cell types by assessing the concordance between TF expression and the trait regulation score.

In our specific study on inflammatory bowel disease (IBD), we collected the risk loci based on the summary statistics of a GWAS meta-analysis of about 330,000 individuals from the NHGRI-EBI GWAS Catalog [51] for study GCST90225550 [52]. Fig. 5A shows that in classical monocyte cells, trait regulation scores for the top 100 expressed TF are significantly higher than randomly selected TFs (P=8.9e-26, one side two sample t-test), while there is no significant difference between top and random selected TF in trait non-relevant cell Types like CD56 (dim) NK cells (P=0.25, one sided unpaired t-test). Across all cell types in PBMCs, the most relevant cell types are classical, non-classical, and intermediate monocytes, as well as myeloid DC (Fig. 5B). These findings align with previous studies linking monocytes to the pathogenesis of IBD [53,54].

**Fig. 5.**
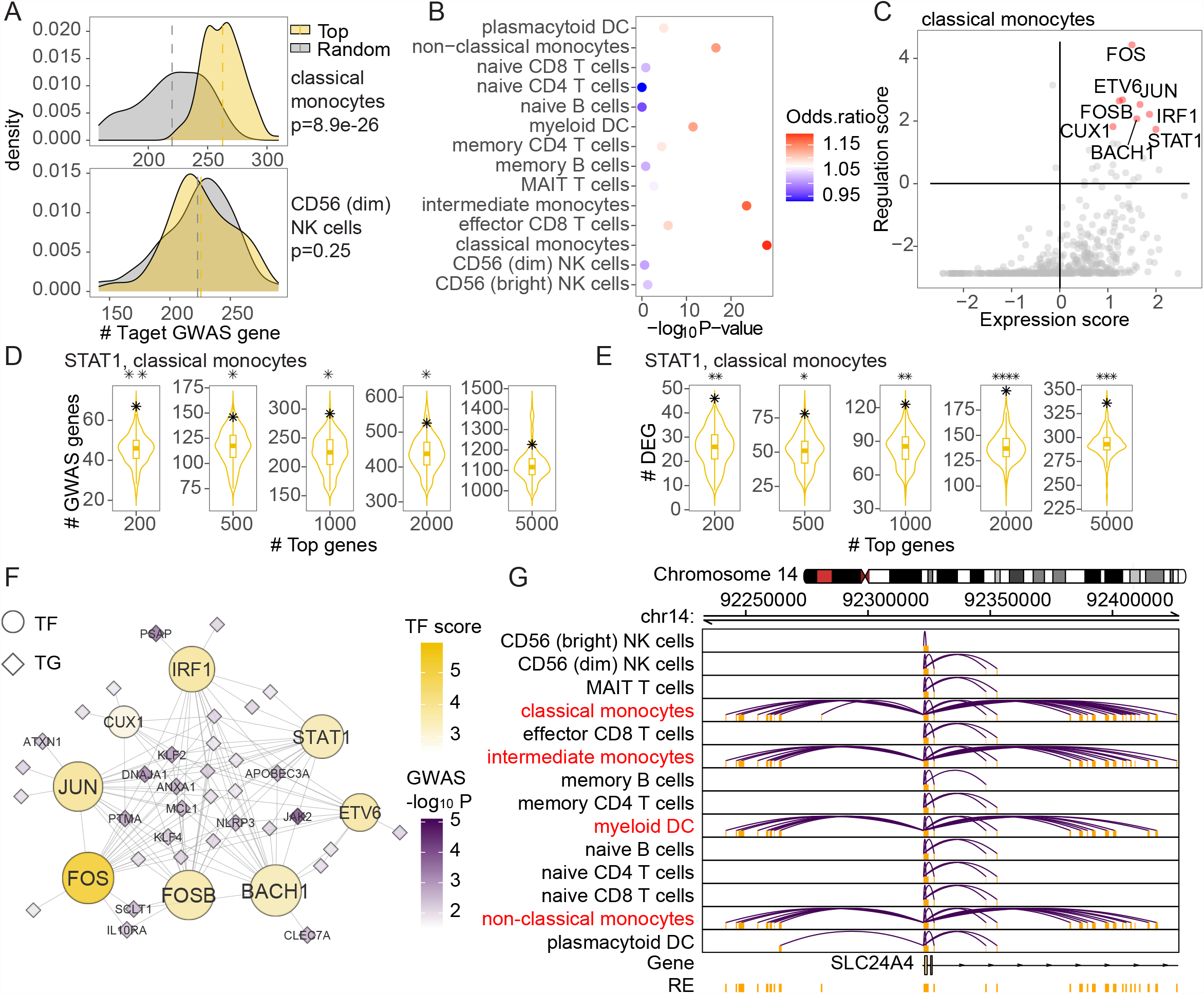
Elucidating GWAS traits through LINGER inferred regulatory landscape. **A**. Distribution of the number of target genes for top expression TFs and randomly selected TFs in classical monocytes (top) and CD56 (dim) NK cells (bottom) 100 top expression TFs and the 100 randomly selected TFs are employed to generate the distribution. **B**. Enrichment of IBD GWAS to cell types in PBMCs. The color of the bubbles corresponds to the odds ratio of the number of target genes between top expression and randomly selected TFs. The x-axis is the -log_10_ P-value from the one sided unpaired t-test for the number of target genes between top expression and randomly selected TFs. **C**. Key IBD-associated regulators in classical monocytes. The x-axis is the z-score of the expression of TFs across all TFs. The y-axis is the regulation score of TFs. The TFs in red color are the top-ranked TFs according to the summation of the expression level and regulation score. **D**. Enrichment of GWAS IBD genes among STAT1 targets in classical monocytes. The violin plot is generated by randomly choosing 1000 TFs and the number of overlapping genes for STAT1 is marked by star. The different violin plots correspond to taking the top 200 to 5000 genes as the target gene for each TF, respectively. **E**. Enrichment of differentially expressed genes between inflamed biopsies and non-inflamed biopsies among STAT1 targets in classical monocytes. The detail is the same with D. **F**. Sub-network of IBD-relevant TFs from classical monocytes *trans*-regulatory network. The round node represents TF, while the rhombus represents TG. The size of nodes corresponds to the degree. The color of TF denotes the trait-relevant score, and the color of TG denotes the - log_10_ P-value of GWAS SNP assigned to the gene. **G**. *Cis*-regulatory network at locus around SLC24A4. The interaction denotes significant RE-TG links, and we use the location of the promoter to represent the gene.

We next identified key TFs by defining the trait-relevant score as the summation of the expression level and trait regulation score for prioritized cell types. Fig. 5C shows the most relevant TFs in classical monocytes and list the top eight candidate TFs. These TFs have been previously reported to be associated with IBD in literature. For example, FOS could increase the risk of recurrence of IBD [55]; One variant identified in IBD cohort is located at the exon of ETV6; IRF1 and ETV6 are key TFs with activity differences in IBD [56]; FOS, FOSB, and JUN are potent mediators of IBD [57]; CUX1 is induced in inflammatory bowel disease [58]; STAT1 epigenetically contribute to the pathogenesis of IBD [59].

To investigate the downstream targets of key TFs, we chose STAT1 as an example to illustrate how GRN bridges GWAS and differential expression analysis. STAT1 is one of the candidate TFs in classical monocytes. Among the top 200 TGs regulated by STAT1 in classical monocytes, 67 of them overlap with the GWAS genes. To determine the significance of this overlap, we perform a random selection of TFs to generate a background distribution of the number of overlapped genes. The results indicate that the observed overlap of 67 genes is statistically significant with a p-value of less than 0.01. We also did the same analysis for the top 500, 1000, 2000, and 5000 TGs, and the results are all significant (Fig. 5D). Apart from GWAS relevant gene, we collected the differentially expressed genes (DEGs) between inflamed biopsies and non-inflamed biopsies [60], and we found these DEGs are significantly overlapped with the top-ranked TGs of STAT1 (Fig. 5E). The lack of significant overlap between DEGs and GWAS genes (P= 0.15, Fisher’s exact test), but the significant overlap of both DEGs and GWAS with the top-ranked TGs of STAT1, indicates the robustness and unbiased nature of our method.

Finally, we extracted the sub-network of the eight candidate TFs from the classical monocyte *trans*-regulatory network, which represents the most relevant TFs and target genes for IBD (Fig5. F). This network shows that many GWAS genes are regulated by multiple key TFs in classical monocytes. We also observed that the *cis*-regulatory network of SLC24A4 (Fig5. G), 46kb from a risk SNP rs11626366 (p= 7.4e-3), is specifically dense in the IBD relevant cell types, which shows the complex regulatory landscape of disease genes across different cell types.

### Identify driver regulators based on transcription profiles

Researchers commonly utilize bulk or single-cell gene expression data from cases and controls to identify DEGs. However, a critical question remains: What are the underlying driver regulators responsible for these observed changes? TF activity, focusing on the DNA-binding component of TF proteins in the nucleus, is a more reliable metric than mRNA or whole protein expression for identifying driver regulators. Direct measurement of TF activity can be challenging. However, one feasible approach is to estimate TF activity based on the expression patterns of downstream target genes, which necessitates the availability of an accurate GRN. Here, we employed LINGER inferred GRNs from sc-multiome data of a single individual. Assuming the GRN structure is consistent across individuals, we estimated TF activity using gene expression data alone. By comparing TF activity between cases and controls, we identified driver regulators. This approach is valuable as it leverages limited sc-multiome data to estimate TF activity in multiple individuals only using gene expression data (See Methods). To demonstrate the effectiveness of our approach, we present two illustrative examples showcasing its utility in identifying driver regulators.

Example 1: We collected the bulk gene expression data from 26 acute myeloid leukemia (AML) patients and 38 healthy donors [61]. After calculating the TF activity based on the cell population GRN for these samples, we found that FOXN1 is significantly less active in AML patients compared to healthy donors (P=0.035, two tail unpaired t-test; Fig. 6A), while it is not differentially expressed (P = 0.23, two tail unpaired t-test; Fig. 6B). In addition, we calculated the TF activity based on the transcriptome profile of 671 AML individuals [62] and performed survival analysis, which [62] shows that individuals with high FOXN1 activity level tend to have a higher survival probability (Fig. 6C). Furthermore, FOXN1 has been reported as a tumor suppressor [63].

**Fig. 6.**
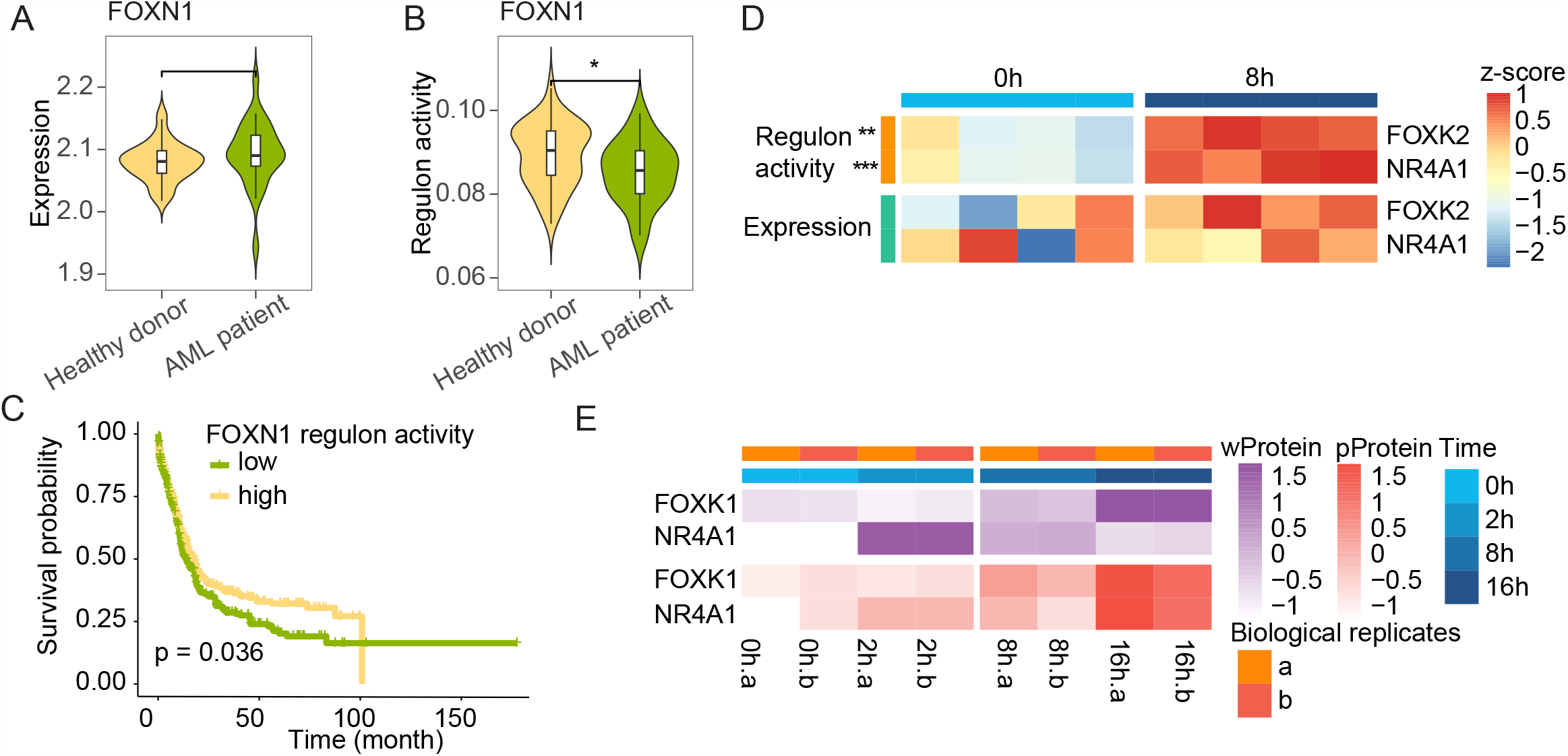
Driver regulator identification. **A**. Violin plot of FOXN1 expression across healthy donors and AML patients, respectively. There is no significant difference in the mean expression under the two sided unpaired t-test. **B**. Violin plot of regulon activity of FOXN1 across healthy donors and AML patients, respectively. **C**. Acute Myeloid Leukemia survival by the regulon activity of FOXN1. **D**. The heatmap of regulon activity and gene expression in response to TCR stimulation at 0h and 8h. The two sided unpaired t-test for the difference in regulon activity gives the p-value of 0.0057 and 0.00081 for FOXK1 and NR4A1, respectively, while the p-value on gene expression is bigger than 0.05. The heatmap is scaled by row. **E**. The heatmap of whole protein and phosphoproteomics expression in response to TCR stimulation at 0h, 2h, 8h, and 16h. There are 2 biological replicates represented by a and b. The whole protein and phosphoproteomics expression of FOXK1 and NR4A1 are higher in 8h compared with 0h. The heatmap is scaled by row.

Example 2: We also present an example of capturing driver regulators of the naive CD4+ T-cell response upon T-cell receptor (TCR) stimulation [63], which induces T cell differentiation into various effector cells and activates T lymphocytes. We calculated the TF activity based on the GRN of naïve CD4+ T cells and identified differentially active regulators in response to TCR stimulation at 8 h vs. 0 h. FOXK2 and NR4A1 are activated at 8h based on regulon activity (Fig. 6D), which is consistent with the whole proteomics (wProtein) and phosphoproteomics (pProtein) data (Fig. 6E) [64]. Other studies have also shown that FOXK2 affects the activation of T lymphocytes [65,66] and revealed the essential roles of the NR4A1 in regulatory T cell differentiation [67,68], suggesting the identified TFs play important roles in naive CD4+ T-cell response on TCR stimulation. However, FOXK2 and NR4A1 show no significant differences in expression at 8 h vs. 0 h (Fig. 6D).

## Conclusions and discussions

LINGER is a deep learning-based method for inferring GRNs from single-cell multiome data by incorporating external data and knowledge of TF-RE motif matching to address the challenge of limited data and a large parameter space in GRN inference. Our results demonstrate that LINGER achieves much higher accuracy compared to existing methods, and it also allows for the estimation of TF activity solely from bulk or single cell gene expression data, to leverage the abundance of available gene expression data to identify driver regulators from case-control studies.

Discussion 1: Computational methods for GRN inference have a long history, but a major challenge has been the limited availability of data and the vast parameter space, making it difficult to accurately fit complex models. In this context, lifelong machine learning emerges as an ideal technology, leveraging external bulk data for diverse tissues, paired single-cell data for numerous cells, and increased computational power. By continuously updating the model with new data, lifelong learning can effectively adapt and improve GRN inference performance, enabling more accurate and comprehensive insights into gene regulatory interactions. The Lifelong learning approach in LINGER has the added advantage of being pre-trained on diverse cellular contexts, allowing users to easily retrain the model with their own data to effectively leverage the wealth of publicly available data without necessitating direct access to that data.

Discussion 2: Traditionally, the accuracy of a regression-based GRN inference method in predicting gene expression has been considered an indicator of its ability to capture underlying regulatory mechanisms and yield improved GRN inference. However, the use of lifelong learning to leverage external data doesn’t necessarily lead to improved gene expression prediction task performance. Nevertheless, it significantly improves the GRN inference task. This finding challenges the traditional strategy of evaluating GRN inference solely based on gene expression prediction and highlights the importance of considering other factors, such as the overall network structure and regulatory interactions, when assessing the performance of GRN inference methods.

Discussion 3: An important feature of the LINGER method is its capacity to infer TF activity using the GRNs obtained from sc-multiome data. By inputting gene expression data from bulk or single-cell samples, researchers can deduce TF activity levels and identify hidden drivers through differential activity testing. This approach is valuable as it seamlessly integrates sc-multiome data with widely available gene expression data, enabling its application in various studies. The simplicity of inferring TF activity from readily available gene expression data makes it accessible to researchers, facilitating the exploration of regulatory interactions within specific biological and clinical contexts. This integration of GRN inference and TF activity prediction in LINGER holds great promise, revolutionizing our understanding of regulatory networks and their implications in diverse biomedical systems.

## Methods

### GRN inference by lifelong learning

LINGER is a computational framework to infer gene regulatory networks (GRNs)—pairwise regulation among target gene (TG), regulatory element (RE), and transcription factor (TF)—from single-cell multiome data. Overall, LINGER predicts the gene expression by the TF expression and chromatin accessibility of REs based on neural network (NN) models. The contribution of each feature is estimated by the Shapley value of the NN models, enabling the inference of the GRNs. To capture key information from the majority of tissue lineages, LINGER utilizes lifelong machine learning (continuous learning). Moreover, LINGER integrates motif binding data by incorporating a manifold regularization into the loss function.

The inputs for full training of LINGER are external bulk and single-cell paired gene expression and chromatin accessibility data. However, we provided a bulk data pretrained LINGER model so that user can retrain it for their own single cell data without accessing to external bulk data. We collect paired bulk data—gene expression profiles and chromatin accessibility matrices—from 201 samples from diverse cellular contexts [69] from ENCODE project [70]. Single-cell data are raw count matrices of multiome single-cell data (gene counts for RNA-seq and RE counts for ATAC-seq). LINGER trains individual models for each gene using a neural network architecture that includes a single input layer and two fully connected hidden layers. The input layer has dimension equal to the number of features, containing all TFs and REs within 1Mb from the transcription start site (TSS) for the gene to be predicted. The first hidden layer has 64 neurons with rectified linear unit (ReLU) activation, which could capture regulatory modules each of which contain multiple TFs and REs. These regulatory modules are characterized by enriched motifs of the TFs on the corresponding REs. The second hidden layer has 16 neurons with ReLU activation. The output layer is a single neuron, which outputs a real value for gene expression prediction.

We first construct NN models based on bulk data, using the same architecture described above. We extract the TF expression matrix 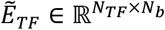 from the bulk gene expression matrix 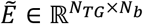, with *N*_*TG*_ representing the number of genes, *N*_*TF*_ representing the number of TF, and *N*_*b*_ representing the number of tissues. The loss function consists of mean squared error (MSE) and L1 regularization, and here is the loss function of the *i* th gene:

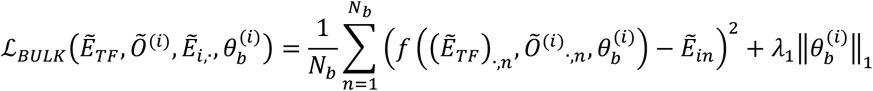

Where 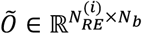 represents the chromatin accessibility matrix, with 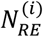 REs within 1Mb from the TSS of the *i* th gene; 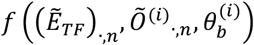 is the predicted gene expression from the neural network of the sample *n*. The neural network is parametrized by a set of weights and biases, collectively denoted by 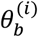. The weight *λ*_1_ is a tuning parameter.

The loss function of LINGER is composed of MSE, L1 regularization, manifold regularization, and elastic weight consolidation (EWC) loss, which is *ℒ*_*LINGER*_ = *λ*_1_*ℒ*_*MSE*_ + *λ*_2_*ℒ*_*L*1_ + *λ*_3_*ℒ*_*Laplace*_ + *λ*_4_*ℒ*_*EWC*_. *ℒ*_*Laplace*_ represents the manifold regularization because a Laplacian matrix is utilized to generate this regularization term. The loss function terms correspond to the gene *i*, and for simplicity, we omit subscripts—(*i*)—for the chromatin accessibility matrix (*O*), parameter for the bulk model (*θ*_*b*_), and parameter for LINGER (*θ*_*l*_).

1. mean squared error (MSE).

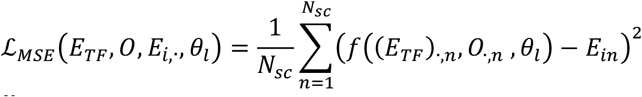

Where 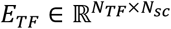 represents the TF expression matrix from the single-cell RNA-seq data, consisting of *Nsc* cells; 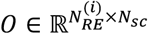 represents the RE chromatin accessibility matrix of the single-cell ATAC-seq data; 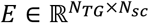 represents the expression of the genes across cells; *θ*_*l*_ represents the parameters in the neural network. Here, we use pseudo-bulk to train the models, therefore *N*_*sc*_ is the number of cells from pseudo-bulk data.
2. L1 regularization.

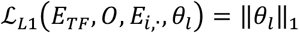
3. Laplacian loss (manifold regularization). We generate the TF-RE adjacency matrix *B* * ∈ 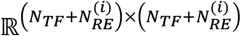, where 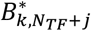 and 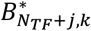 represent the binding affinity of the TF *k* and the RE *j*., respectively 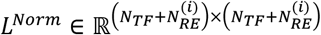 is normalized Laplacian matrix, which is elaborated in the following sections.

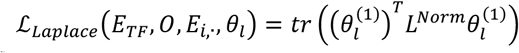

Where 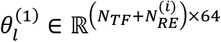 is the parameter matrix of the first hidden layer, which could capture the densely connected TF-RE modules.
4. EWC loss. EWC constrains the parameters of the first layer to stay in a region of 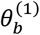, which are previously learned from bulk data [36]. To do this, EWC uses MSE between the parameters 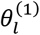 and 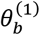, weighted by the Fisher information, a metric of how important the parameter is, allowing for the model protecting the performance both for single-cell data and bulk data [36].

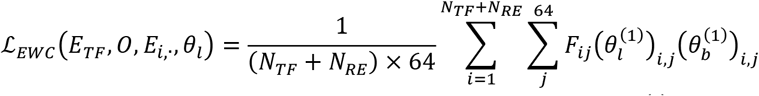

Where *F* is the fisher information matrix, which is detailed below; 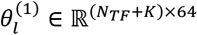 is the parameter matrix of the first hidden layer.

To construct a normalized Laplacian matrix, we first generate the TF-RE binding affinity matrix for all REs from the single-cell ATAC-seq data. We extract the REs 1Mb from the TSS for the gene to be predicted. Let 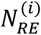 be the number of these REs and 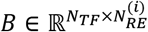 be the TF-RE binding affinity matrix, where *B*_*kj*_ represents the binding affinity for the TF *k* and RE *j*. We construct a graph, taking TFs as the first *N*_*TF*_ nodes, REs as the remaining 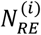 nodes, and binding affinity as edge weight between TF and RE. The edge weights of TF-TF and RE-RE are set to zero. Then adjacency matrix 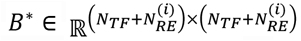 is defined as:

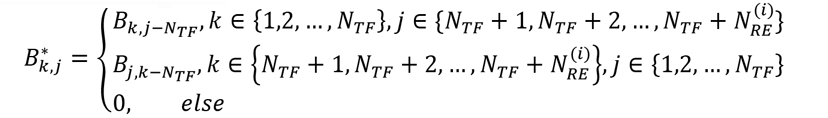

The Fisher information matrix is calculated based on the neural network trained on bulk data,

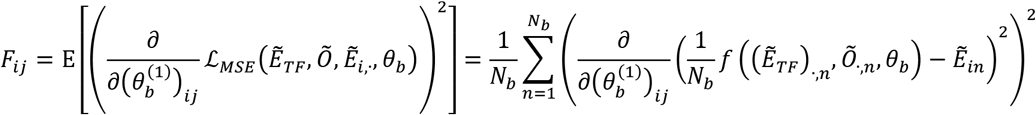

### GRN inference by Shapley value

Shapley value could measure the contribution of features in a machine learning model, which is widely used to explain the models, such as deep learning, graphical models, and reinforcement learning [71]. We use the average of absolute Shapley value across samples to infer the regulation strength of TF and RE to target genes, generating the RE-TG *cis*-regulatory strength and the TF-TG *trans*-regulatory strength. Let *β*_*ij*_ represents the *cis*-regulatory strength of RE *j* and TG *i*, and *γ*_*ki*_ represents the *trans*-regulatory strength. To generate the TF-RE binding strength, we use the weights from input layer (TFs and REs) to all node in the second layer of the NN model as the embedding of the TF or RE. The TF-RE binding strength is calculated by the PCC between the TF and RE based on the embedding. *α*_*kj*_ represents the TF-RE binding strength.

### Construct cell type-specific gene regulatory network

TF-RE regulatory potential for a certain cell type is given by,

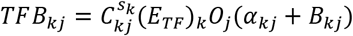

Where *TFB*_*kj*_ is the TF-RE regulation potential of TF *k* and RE *j*; *s*_*k*_ is a importance score of TF *k* in the cell type to measure the preference of TF for activating cell type-specific open chromatin regions, which will be described in section “TF importance score”; *C*_*kj*_ is the PCC of TF *k* and RE *j*; *O*_*j*_ is the average chromatin accessibility across cells in the cell type; *B*_*kj*_ is the binding affinity between TF *k* and RE *j*; *α*_*kj*_ is the TF-RE binding strength.

RE-TG *cis*-regulatory potential is defined as:

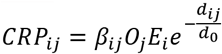

Where *CRP*_*ij*_ is the *cis*-regulatory potential of TG *i* and RE *j*; *β*_*ij*_ is the *cis*-regulatory strength of RE *j* and TG *i*; *O*_*j*_ is the average chromatin accessibility; *E*_*i*_ is the average gene expression across cells in the cell type; *d*_*ij*_ is the distance between genomic locations of TG *i* and RE *j*; *d*_0_ is a fixed value used to scale the distance, which is set to 25k in this paper.

TF-TG *trans*-regulatory potential is defined as the cumulative effect of corresponding REs on the TG:

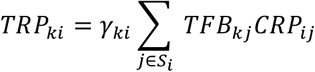

Where *γ*_*ki*_ is the TF-TG *trans*-regulatory strength of TF *k* and TG *i*; *S*_*i*_ is the set of RE within 1Mb from TSS for TG *i*; *CRP*_*ij*_ is the *cis*-regulatory potential of TG *i* and RE *j*; *TFB*_*kj*_ is the TF-RE regulation potential of TF *k* and RE *j*.

### Construct cell-level GRNs

Cell-level GRNs are inferred by integrating information consistent across all cells, such as regulatory strength, binding affinity, and RE-TG distance, with cell-level information, such as the gene expression and chromatin accessibility. This approach is similar to inferring cell type specific GRN with the key difference that cell-level GRN use cell-level TF expression *E*_*TF*_, chromatin accessibility *O*, and gene expression *E* rather than cell type-averaged data. This allows us to infer the network for each individual cell based on its specific characteristics, rather than grouping cells into predefined types.

### TF importance score

To systematically identify TFs playing a pivotal role in controlling the chromatin accessibility of cell type, we introduce a TF importance score. The score is designed to measure the preference of TF for activating cell type-specific REs. The input is multiome single-cell data with known cell type annotations. There are steps to generate the TF importance score:

1. Motif enrichment. We perform the motif enrichment analysis [72] to identify the motifs significantly enriched in the binding sites of the top 5,000 cell type-specific REs. We use the p-value to measure the significant level of motif enrichment.
2. TF-RE correlation. To avoid dropouts in single-cell data, we recover the original count matrix by an average of the observed count of nearby cells. We calculate Pearson’s correlation coefficient (PCC) between the TF expression and cell type specific RE chromatin accessibility with *r*_*kj*_ representing the PCC of the TF *k* and the RE *j*. To mitigate the bias in the distribution of TF expression and REs chromatin accessibility so that the PCC is comparable across different TF-RE pairs, we permute the cell barcode in the gene expression data and then calculate, generating a background PCC distribution for each TF-RE pair. We generate a z-score for *r*_*kj*_,

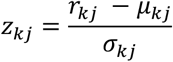

Where *μ*_*kj*_ and 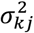 is the mean and the variance of background PCC distribution between *TF*_*k*_ and *RE*_*j*_,
3. The co-activity score of the TF-motif pair. To pair TFs with their motifs, we match 713 TFs and 1,331 motifs, yielding 8,793 TF-motif pairs [69]. Let (*k, m*) denotes the TF-motif pair of TF *k* and motif *m*. We then calculate a co-activity score for a TF-motif pair for (*k, m*), defined as the average z-score across cell type-specific REs with at least one motif binding site. That is 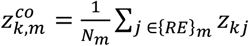, where {*RE*}_*m*_ is the set of REs with the *m*-th motif binding; *N*_*m*_ = |{*RE*}_*m*_| is the number of RE in {*RE*}_*m*_.
4. TF importance score. The score of the TF-motif pair, (*k, m*), is given by,

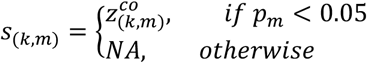

Where *p*_*m*_ is the p-value of the *m* -th motif from the motif enrichment analysis; *s*_(*k,m*)_ is the importance score of the TF-motif pair (*k, m*). The TF importance score for the TF *k* is the average TF-motif pair TF importance score across motifs, omitting NA:

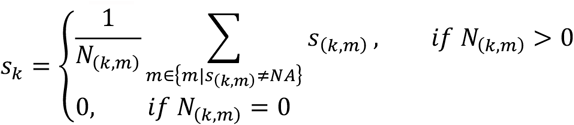

Where *N*_(*k,m*)_ = |{*m*|*s*_(*k,m*)_ ≠ *NA*}| is the number of the TF-motif pair of the TF *k*, whose CECI score is not NA.

### TF-RE binding affinity matrix

We download 713 TF position weight matrices (PWMs) for the known motifs from GitHub page of PECA2 [69], which is collected from widely used databases, including JASPAR, TRANSFAC, UniPROBE, and Taipale. Given a list of REs, we calculate the binding affinity score for each TF by motif scan using Homer [72], as a quantitative measure of the strength of the interaction between TF and RE [15].

### Identify motif-binding regulatory elements

We identify the REs with motif binding by motif scan using Homer [72].

### PBMC 10x data

We download the PBMC 10K data from the 10X genomics website https://support.10xgenomics.com/single-cell-multiome-atac-gex/datasets. Note that it contains 11,909 cells, and the granulocytes were removed by cell sorting of this dataset. We use the filtered cells by features matrix from the output of 10X genomics software Cell Ranger ARC as input and perform the downstream analysis. First, we perform Seurat 4.0 [37] weighted nearest neighbor (WNN) analysis and it removes 1,497 cells. We also remove the cells that do not have surrogate ground truth and it results in 9,543 cells. We generated pseudobulk data by randomly selecting square root of the number of cells in each cell type and averaged the expression levels and chromatin accessibility of 100 nearest cell to produce the gene expression and chromatin accessibility values of the selected cells. The pseudobulk data were directly input into LINGER for analysis.

### AUPR ratio

To measure the accuracy of a predictor, we defined the area under the precision-recall curve (AUPR) ratio as the ratio of the AUPR of a method to that of a random predictor. For a random predictor, the AUPR equals the fraction of positive samples in the dataset. The AUPR ratio is defined as the 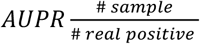, representing the fold change of the accuracy of a predictor compared to the random prediction.

### LINGER reveals the complex regulatory landscape of GWAS traits

We propose a method to integrate GWAS summary statistic data and cell type specific GRNs to identify the relevant cell types, key TFs, and sub-GRN responsible for GWAS variants. To identify relevant cell types, we first project the risk single nucleotide polymorphism (SNP) identified from GWAS summary data to gene. We then link gene within the 200 kb region centering at the SNP and assign the most significant p-value of linked SNPs to each gene. In this study, the trait related gene are defined as those with a p-value <0.01 after FDR adjustment. We then calculate a trait regulation score for each TF in each cell type, measuring the enrichment of GWAS genes downstream of the TF based on the cell type specific GRN. We choose 1000 top-ranked gene according to the *trans*-regulation as the target gene of each TF and count the number of overlapping genes with trait-related genes. The enrichment of cell types to the GWAS traits is measured by a t-test comparing the number of the overlap gene between the 100 top expressed and 100 randomly chosen TFs.

To identify key TFs of GWAS traits, we combine the trait regulation score and the gene expression level of TFs in each cell type. The trait regulation score is the z-score of the number of overlapping genes of a TF across all TFs. The expression level is also transformed to a z-score based the gene expression. The final importance of key TFs is the summation of the expression level and trait regulation score.

### Identify driver regulators based on transcription profiles

To measure the activity of each TF on the independent transcriptional profiles, we first constructed a target gene set for each TF based on the corresponding GRN. We perform quantile normalization to the trans-regulation score of each gene across all TFs. We then rank the genes for each TF and chose top 1000 genes as the target. Next, we use the R package AUCell [17] to calculate whether the target genes are enriched within the expressed genes for each sample, which defines the TF activity.

### Benchmark the *trans*-regulatory potential

We compare LINGER’s performance of the trans-regulation prediction using PCC, SCENIC+, GENIE3, and PIDC as competitor to LINGER. Due to the time-consuming nature of PIDC’s mutual information-based algorithm, we used the 5000 most variable genes as input. As a result, there are 9 TFs and 14 TFs in ground truth data left for PBMCs and H1 cell line, respectively.

## Data availability

The PBMCs data used during this study is downloaded from the 10X Genomics website [33]. SNARE-seq is downloaded from NCBI Gene Expression Omnibus (GEO, https://www.ncbi.nlm.nih.gov/geo/) under accession number GSE126074 [45].

## Code availability

The software is available at GitHub at https://github.com/Durenlab/LINGER.

## Acknowledgements

This work was partially supported by NIH grants P20 GM139769.

## Author contributions

Z.D. conceived the LINGER method. Z.D. and Q.Y. designed the analytical approach. Q.Y. performed the data analysis. Q.Y. wrote the software. Q.Y. and Z.D. wrote, revised, and contributed to the final manuscript. The authors read and approved the final manuscript.

