## Supplementary material for "Continuous lifelong learning for modeling of gene regulation from single cell multiome data by leveraging atlas-scale external data": Fig. S1 and Fig. S2

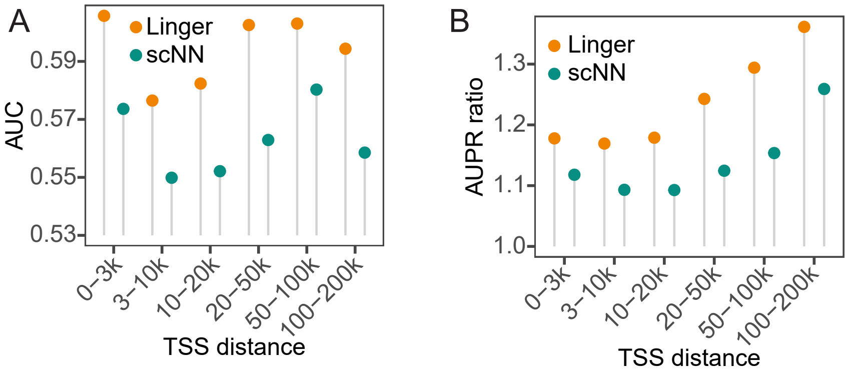


**Fig. S1|** Assessing the performance of *cis*-regulatory strength inferred by LINGER taking eQTL data for GTEx as ground truth. **A**. AUC for *cis*-regulatory strength inferred by LINGER. The ground truth for A and B is the variant-gene links from GTEx. We divide RE-TG pairs into different groups based on the distance of RE and the TSS of TG. **E**. AUPR ratio for *cis*-regulatory strength.

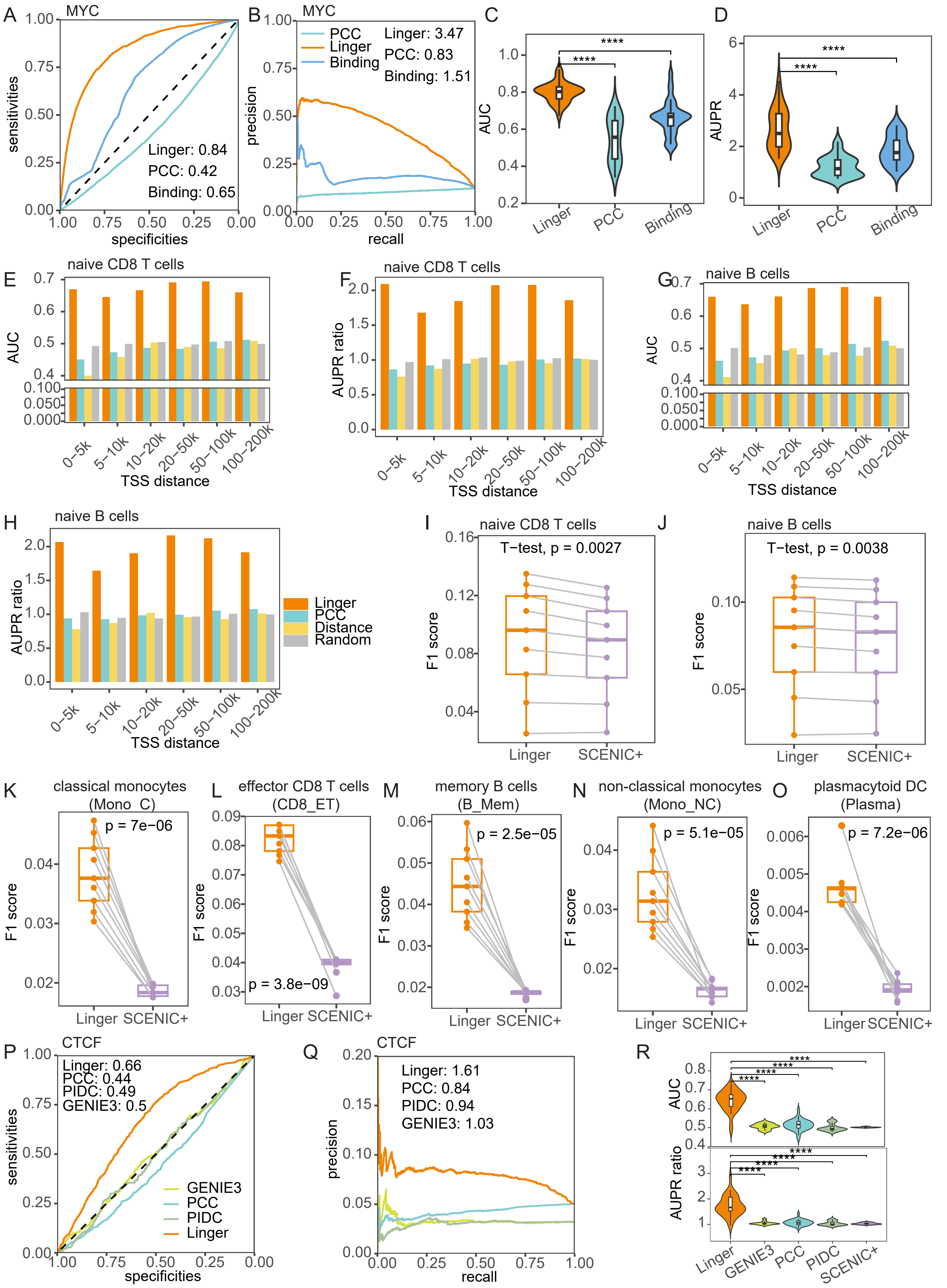

A
MYC
sensitivities
specificities
Linger: 0.84
PCC: 0.42
Binding: 0.65
B
MYC
precision
recall
PCC
Linger
Binding
Linger: 3.47
PCC: 0.83
Binding: 1.51
C
AUC
****
Linger
PCC
Binding
D
AUPR
****
Linger
PCC
Binding
E
naive CD8 T cells
AUC
TSS distance
0–5k
5–10k
10–20k
20–50k
50–100k
100–200k
F
naive CD8 T cells
AUPR ratio
TSS distance
G
naive B cells
AUC
TSS distance
H
naive B cells
AUPR ratio
TSS distance
Linger
PCC
Distance
Random
I
naive CD8 T cells
T−test, p = 0.0027
F1 score
Linger
SCENIC+
J
naive B cells
T−test, p = 0.0038
F1 score
Linger
SCENIC+
K
classical monocytes
(Mono_C)
p = 7e−06
F1 score
Linger SCENIC+
L
effector CD8 T cells
(CD8_ET)
p = 3.8e−09
F1 score
Linger SCENIC+
M
memory B cells
(B_Mem)
p = 2.5e−05
F1 score
Linger SCENIC+
N
non-classical monocytes
(Mono_NC)
p = 5.1e−05
F1 score
Linger SCENIC+
O
plasmacytoid DC
(Plasma)
p = 7.2e−06
F1 score
Linger SCENIC+
P
CTCF
Linger: 0.66
PCC: 0.44
PIDC: 0.49
GENIE3: 0.5
sensitivities
specificities
GENIE3
PCC
PIDC
Linger
Q
CTCF
Linger: 1.61
PCC: 0.84
PIDC: 0.94
GENIE3: 1.03
precision
recall
R
AUC
AUPR ratio
****
Linger
GENIE3
PCC
PIDC
SCENIC+

**Fig. S2|** Systematic benchmarking of cell type specific GRN. **A, B**. ROC curve and PR curve of binding potential for MYC in H1 cell line. The ground truth for A to D is the ChIP-seq data of MYC in the H1 cell line. The color in A to D represents the different competitors to predict TF-RE regulation. Orange represents LINGER, green represents PCC between the expression of TF and the chromatin accessibility of RE, and blue represents motif binding affinity of TF to RE. **C, D**. Violin plot of AUC and AUPR ratio values of binding potential across diverse TFs. The ground truth is ChIP-seq data for 33 TFs. **E, F**. AUC and AUPR ratio of *cis*-regulatory potential in naïve CD8 T cells. The ground truth for I to J is promoter capture HiC data. RE-TG pairs are divided into six distance groups ranging from 0-5k to 100-200 kb. PCC is calculated between the expression of TG and the chromatin accessibility of RE. Distance denotes the decay function of the distance to the TSS. Random denotes the uniform distribution. **G, H**. AUC and AUPR ratio of *cis*-regulatory potential in naïve B cells. **I**, **J**. F1 score of *cis*-regulatory in naïve CD8 T cells and naïve B cells for LINGER and SCEINC+. **K** to **O**, F1 score of *cis*-regulatory potential in classical monocytes, effector CD8 T cells, memory B cells, non-classical monocytes, and plasmacytoid DC cells for LINGER and SCEINC+. The ground truth is eQTL data. **P, Q**. ROC curve and PR curve of *trans*-regulatory potential inference of CTCF in H1 cell line. The ground truth of P to R is putative targets of TFs from ChIP-seq data in the H1 cell line. **R** Violin plot of AUC and AUPR ratio values of *trans*-regulatory potential performance across diverse TFs in H1 cell line.
